# Expression of photoactivatable molecules enables FCS in live cells by controlling fluorescence intensity

**DOI:** 10.64898/2026.08.11.743982

**Authors:** Niaz Z. Goodbee, Don Teasley, Christian Pagán Medina, Mary W. Elting, Sharonda J. LeBlanc

**Affiliations:** Department of Physics and Astronomy, North Carolina State University, Raleigh, NC 27695-8202, USA; Microbiology Graduate Program, North Carolina State University

**Keywords:** Fluorescence correlation spectroscopy (FCS), photoswitchable fluorescent protein, nucleoplasm material properties, nuclear mechanics, *Schizosaccharomyces pombe*

## Abstract

Cellular systems must act robustly to maintain organismal health, including maintaining biophysical properties that allow for appropriate cellular function, and adapting these properties through changes such as those that occur during cell division. However, we still lack tools to measure many of these physical properties with precision in the living cell. For example, the mechanical properties of the nucleoplasm, the fluid-like substance that fills the nucleus, have not been fully characterized. To investigate these properties, we have turned to the fission yeast *Schizosaccharomyces pombe* (*S. pombe*), a well-established, genetically tractable model organism that has been used extensively for studying a variety of cell biophysical processes and structures, including the cytoskeleton and cell division. It is an apt system for studying how the nucleus adapts over the course of the cell cycle, since it undergoes closed mitosis, where the nuclear envelope remains intact during cell division. Studying nucleoplasm properties over the course of closed mitosis may help reveal how nuclear volume, shape, surface area expansion, and chromosome segregation are linked and coordinated. To measure nucleoplasm material properties in *S. pombe*, we have paired Fluorescence Correlation Spectroscopy (FCS) with a photoswitchable fluorophore, enabling fine control over fluorescent intensity inside live cells. We infer material properties from FCS measurements, while the photoswitchable probe enables confocal imaging in conjunction with these measurements, yielding corresponding information about cellular state and dynamics. Interestingly, we find that nucleoplasm material properties do not vary significantly over the cell cycle. Future studies will use this tool to examine how diverse molecular and genetic perturbations alter nucleoplasmic properties, providing insight into how these properties maintain nuclear function and protect genomic integrity over the cell cycle and during development.

## Introduction

Cellular changes, whether from osmotic shifts, cytoskeletal forces, or alterations in nuclear architecture, can impact cellular material properties, thus altering diffusion, biochemical rates, and ultimately cell function. Understanding the material properties of the nucleoplasm and cytoplasm is therefore essential to physically explaining many biological processes, but is difficult to probe directly. Though both the cytoplasm and the nucleoplasm are mostly water, they are sufficiently crowded with biological molecules to alter their material properties (1–3). Diffusion can be used as a readout of material properties, but comes with its own challenges, since measurements may depend on the size of the probe and other factors such as crowding that influence diffusive behavior (4). Three approaches that have been used to measure diffusion are Fluorescence Recovery After Photobleaching (5, 6), Single Particle Tracking (SPT) (also referred to as Single Molecule Tracking (SMT)) (7–9), and Fluorescence Correlation Spectroscopy (FCS) (10). Each of these three methodologies provides significant analytical power, yet each is accompanied by specific technical constraints.

Developed in the mid 1970s, FRAP came into existence to measure the mobility of molecules but more recently has been used to study binding and active transport (5, 11, 12). This technique works by photobleaching an area or region of interest (ROI) and recording the time it takes to recover the fluorescence from unbleached molecules diffusing into the bleached area (13). In most cases, photobleaching is viewed as deleterious, as it can create cytotoxicity and decreases signal to noise, but FRAP instead leverages it to determine dynamics and kinetics (14, 15). However, this method is limited due to its “ensemble” nature: measuring thousands or millions of molecules can result in missing out on fast diffusing molecules, rare events, and heterogeneous environments (16). FCS and SPT avoid some limitations of FRAP by measuring the trajectory of individual molecules with SPT (17) and capturing diffusion on fast time scales with FCS (18).

Single Particle Tracking (SPT) offers a powerful way to probe the material properties of the cytoplasm and nucleoplasm by analyzing the movement of individual molecules (19). In live cells, SPT can measure diffusion of either fluorescently labeled proteins, organic dyes, or genetically encoded multimeric nanoparticles (GEMs) (20–22). Small molecules diffuse rapidly, making them best suited for SPT when membrane association confines their motion to a 2D surface (7, 8). While 3D tracking in live cells is possible (23, 24), the rapid motion of single molecules can approach the limits of temporal resolution, and their apparent diffusion may be dominated by transient nonspecific interactions (25). A further drawback in three-dimensional samples was highlighted in a study that reported a systematic bias for fast SPT when fitting the mean-squared displacement (MSD) that arises when fast-moving molecules move out of focus and thus lose detectable information (26). In contrast, due to their large size (∼40 nm), GEMs diffuse slowly enough that they can be reliably tracked for longer times. This approach allows diffusive behavior to be extracted from position-over-time measurements, making it a practical way to report on the mechanical response of the cytoplasm and nucleoplasm (20, 27). Experiments with GEMs demonstrated that the nucleus and cytoplasm exhibit distinct material properties, behaving differently in response to changes in crowding and osmotic perturbation (20). The data also suggested that osmotic shifts alone can alter nuclear size and the nuclear-cytoplasmic ratio (N/C), challenging earlier models that attributed nuclear size control primarily to DNA content (28). However, a key limitation of using GEMs to probe material properties is their restricted access: due to their relatively large size, they can only sample regions of the cell with sufficiently large pores. While comparable in size to some biological complexes, such as extracellular lipid vesicles (29) or ribosomes (27), they are much larger than many biomolecules. If the nucleus behaves partly as a poroelastic material, some subnuclear domains may be inaccessible to GEMs, limiting their ability to report on material properties at smaller length scales. Thus, it is important to develop complementary tools that can probe the material properties experienced by more typically-sized biomolecules.

Fluorescence Correlation Spectroscopy (FCS) has been used in many studies (18, 30–32) to measure diffusive behavior by tracking intensity fluctuations as fluorescent molecules diffuse through a confocal volume (33). The autocorrelation of intensity traces allows measurement of diffusion rates, sample concentrations, and molecular interactions. However, a key challenge of FCS is that the detection volume and concentration must be matched: the number of molecules in the confocal observation volume must be low enough, and the signal sampled fast enough, that fluorescence intensity fluctuations can be reliably detected. Because the autocorrelation amplitude decreases as the number of molecules in the detection volume increases, many FCS studies have focused on membrane systems, where molecules are confined to a quasi-two-dimensional geometry and the effective observation volume is reduced by physical confinement relative to the full 3D point spread function (34, 35). Alternative microscopy and spectroscopy implementations, such as rapidly scanning the excitation beam or acquiring fluctuations at multiple measurement points with multiple detectors, extend the spatiotemporal sampling of FCS measurements and enable diffusion mapping in live cells (36, 37). With technological advances in single-photon detectors and multichannel photon-timing instrumentation, FCS has become increasingly sensitive and robust, expanding its feasibility and application across diverse fields in chemistry, biophysics, and cell biology (38).

In this study, we sought to control fluorescence intensity directly, thus making FCS of small molecules in living cells more reliable by not overcrowding the confocal volume. For these studies, we focused on measurements of nucleoplasmic diffusion in closed mitosis, reasoning that the changing structural organization that occurs over cell division might alter material properties in ways that would be difficult to detect via tracking of GEMs. We generated an *S.pombe* strain expressing a photoswitchable fluorescent protein, mEos3.2, tagged to a Nuclear Localization Sequence (NLS) which enables controlling the concentration of activated molecules in the confocal volume. This method provides the further benefit of allowing for quality live cell fluorescence imaging with high signal-to-noise ratio in a spectrally distinct imaging channel. We have used our FCS measurements to compute diffusion rates, and infer nucleoplasm material properties. Through the use of time-resolved fluorescence methods, including Fluorescence Correlation Spectroscopy (FCS) and Fluorescence Lifetime Imaging Microscopy (FLIM), we demonstrated that we can measure 3D diffusion in live nuclei by controlling fluorescence intensity.

## Methods

### Plasmid Design and Strain Construction

Details of all strains are included in Supplementary Table S1. Strains MWE133, MWE144 and MWE145 are new to this work. To generate strain MWE144, a strain expressing mEos3.2-NLS, we used the plasmid toolkit generated by the Martin lab for stable genetic integration (39). More specifically, we began with plasmid pAV0526 (Addgene plasmid # 133500; http://n2t.net/addgene:133500; RRID:Addgene_133500), which integrates into the his5 locus with expression under a thd1 promoter. We modified its design to insert a codon-optimized version of mEos3.2 (sequence obtained from Addgene plasmid # 163100; http://n2t.net/addgene:163100; RRID:Addgene_163100) (40–42) with a C-terminal SV40 Nuclear Localization Signal (NLS). EPOCH Life Sciences (Fort Bend County, Texas, US) then generated the plasmid we designed. Sequence for the codon-optimized mEos3.2 is included in the Supplementary Information (Supplementary Table S1). We then linearized the insertion plasmid and transformed it into parent strain MWE53 (43). We back crossed the transformed strain with the parent and then verified expression of the mEos3.2-NLS gene via fluorescence imaging. This new strain is referenced as MWE144 (see Supplementary Table S1).

For verifying cell stage, it is helpful to have constitutive expression of GFP-labeled tubulin. To generate such a strain (MWE145) expressing mEos3.2-NLS and GFP-atb2, we crossed MWE144 with MWE133 (includes GFP-atb2 expression as well as deletion of Nem1, which we eliminated in the cross, see Table S1) by standard methods. MWE133 in turn was generated by crossing MWE2 (expresses GFP-atb2) and MWE41 (nem1Δ). We verified both of these crossed strains by fluorescence imaging to test for expression of mEos3.2-NLS and GFP-Atb2, and also verified by sequencing the Nem1 locus that an unwanted deletion in parent strain MWE133 was eliminated in the cross. This new strain is referenced as MWE145 (see Supplementary Table S1).

### Sample Prep/Cell Culture

We cultured and prepared *S. pombe* for imaging using standard methods (44). Briefly, we plated cells on YE5S and incubated for 2-3 days at 25℃. For imaging, we picked single colonies and incubated in 2 mL of YE5S media for 12-18 hrs overnight with a rotating drum at 25℃.

After the incubation period, using a spectrophotometer, we measured the optical density at a wavelength of 595 nm. We diluted the sample and incubated for an hour to an OD of 0.2-0.3. For imaging, we prepared gelatin pads of 0.25 mg/mL gelatin in YE5S media on glass coverslips. We then took 1 mL of cells in growth phase, centrifuged them briefly to form a pellet, and removed most of the supernatant. We resuspended the pellet of cells in the remaining media, added 2-5μl of to the gelatin pad, added a coverslip, and sealed with VALAP, (1:1:1 ratio of vaseline:lanolin:paraffin). We identified the cell stage via nuclear morphology and microtubule localization (with the presence of cytoplasmic microtubule bundles indicating interphase and the presence of a mitotic spindle indicating mitosis).

### Time-resolved scanning confocal fluorescence and spinning disc confocal microscopy

For diffusion and fluorescence lifetime imaging microscopy (FLIM) measurements, we used a PicoQuant MicroTime200 confocal laser scanning microscope with a 60x 1.2 numerical aperture (N.A.) water immersion objective as previously described to image cells (45). Picosecond pulsed 485 nm (1.5 - 1.8 kW/cm^2^) and 531 nm (0.6 - 0.72 kW/cm^2^) lasers were used to excite the native and photoswitched mEos3.2 fluorophores, respectively. We used a 405 nm LED (∼900-1000 μW, ThorLabs) to photoswitch mEos3.2. Individual detected photons were spectrally separated between two Single Photon Avalanche Diode (SPAD) detectors into green (Channel 2) and red (Channel 1) emission by dichroic and bandpass optical filters. We used a 511/20 bandpass filter for Channel 2, a 582/64 bandpass filter for Channel 1, and a 532 long pass dichroic mirror to separate the signal. After acquiring initial images, we performed FCS. We positioned the 485 nm or 531 nm laser in the nucleus on at least three different locations with the laser power at 0.054 - 0.255 kW/cm^2^ for 485 nm or 0.02 - 0.03 kW/cm^2^ for 531 nm (Figure 1A,B). Low laser power for the FCS measurements allowed us to acquire multiple measurements without photobleaching the cell or causing phototoxicity.

**Figure 1:**
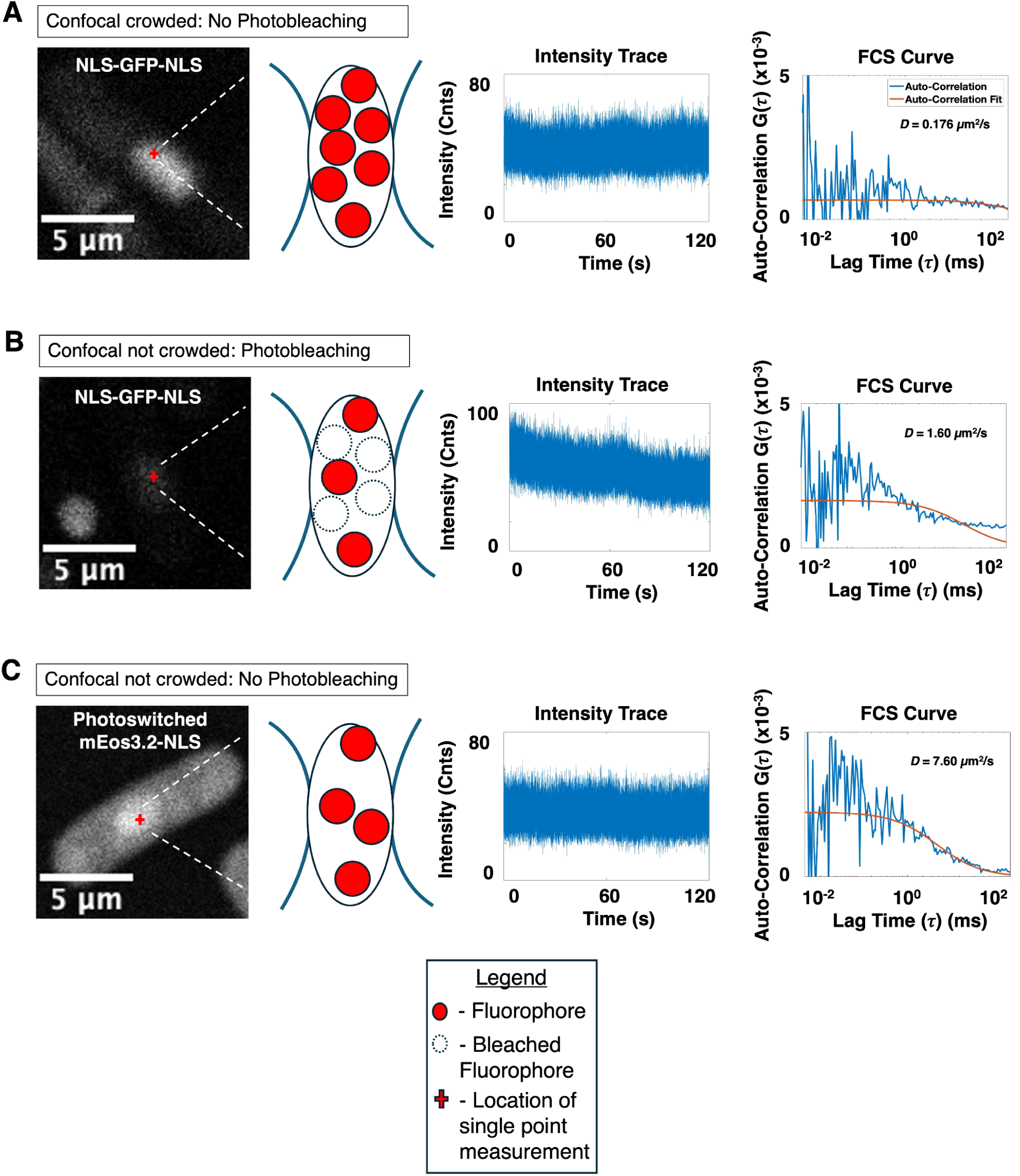
Over-crowding and photobleaching alter the effectiveness of FCS. Each panel shows a representative image, cartoon of the experimental setup, representative intensity trace, and resulting autocorrelation curve and fit. **(A)** Overcrowded confocal volume limits incidence of fluctuations. Single point FCS measurement displays a measured diffusion coefficient of 0.176 μm^2^/s, but is poorly fit due to low signal to noise. **(B)** Photobleaching can improve the incidence of intensity fluctuations for FCS, but results in poor image quality. Single point FCS measurement displays a measured diffusion coefficient of 1.60 μm^2^/s, again with a relatively poor fit. **(C)** An ideal intensity trace and FCS curve resulting in a good fit. Single point FCS measurement displays a measured diffusion coefficient of 7.60 μm^2^/s. Single point FCS measurements collected in the *S. pombe* MWE38 strain expressing NLS-GFP-NLS (A-B) and in *S. pombe* MWE144 strain expressing photoswitched mEos3.2-NLS (C). Scale bars are 5 micrometers across all images.

Andor Dragonfly Spinning Disk Confocal was also used for imaging cells in Figure 3A, as previously described (46). More specifically, a 488 nm (50mW) laser (green channel) and 561nm laser (red channel) were used to excite mEos3.2 and 405 nm LED was used to photoswitch fluorophores. Imaging was captured using a 100x 1.45 Ph3 Nikon oil immersion objective and a 1.5x magnifier (built into the Dragonfly system) along with an Andor iXon3 camera. A spinning disk dichroic (Chroma ZT405/488/561/640rpc) and emission filter (Chroma ET525/50m) was used to separate the signals.

### Fluorescence Correlation Spectroscopy Analysis

Individual detected photons were spectrally separated between the two SPAD detectors into green/native (Channel 2) and red/photoswitched (Channel 1) mEos3.2 emission by dichroic and bandpass optical filters. Fluorescence intensity traces for each channel were generated by grouping sequential photon macrotimes into user-defined equal time bins (usually 1 millisecond). Signal fluctuations were characterized by performing an autocorrelation, *G(τ)*, on the fluorescence intensity signal (Eq. 1):

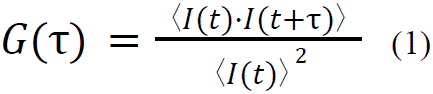

where *I(t)* is the fluorescence intensity over the acquisition time, *τ* is the lag time, and the brackets indicate an average over all time values, *t*. Such a correlation calculates the similarity between the signal and a time-lagged version of itself by changing the lag time, *τ*, in the operation. With increasing *τ*, the time-lagged version becomes less similar to the original signal. We obtained an autocorrelation curve that decays as the lag time increases. Fitting the decay to an appropriate model yields experimental parameters that characterize the physical properties of the fluorescently labeled species (18). To define the diffusion of a small fluorescent molecule, the autocorrelation was fit to a pure diffusion model using Eq. 2:

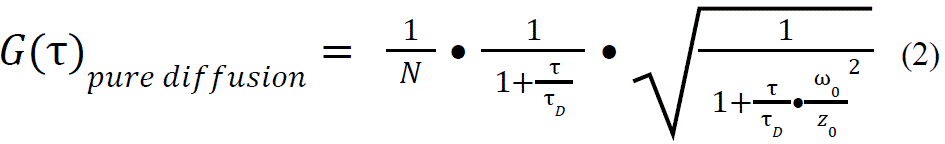

Where the amplitude of the curve is represented by *1/N* at time zero, *N* denotes the average number of molecules in the confocal volume, *τ_D_* is the average time it takes a molecule to diffuse through the confocal volume (translational diffusional time), *ω_0_* is the lateral beam radius, and *z_0_*is the axial beam radius and *z_0_ /ω_0_* is the aspect ratio, *к*. Diffusion coefficients, *D*, were calculated with Eq. 3:

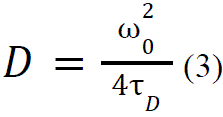

We calibrated the effective confocal volume, *V_eff_* and kappa value, *к* (*z_0_ /ω_0_*) using the dye Rhodamine 6G with a known diffusion coefficient of 400 μm^2^/s (47). For these experiments we have measured the diffusion characteristics of mEos3.2-NLS.

### FLIM Image Analysis

FLIM analysis was performed in SymPhoTime64. Nine neighboring pixels were binned to increase the signal to noise ratio. The Fast Lifetime FLIM image was produced by calculating the average arrival nanotime/microtime for photons in each binned pixel. We selected an ROI from the lifetime and intensity histograms (Supplementary Figure S5(A, B)). The values were set to 3.0 - 5.0 ns for lifetime and 0.8% to 1.3% photons per sync for intensity to reduce contributions from background fluorescence and possible brightspots/aggregates.

## Results

### Transformation and control of activated molecules

FCS, a single-molecule sensitive technique that tracks diffusion by measuring fluorescence intensity fluctuations, requires low fluorophore concentrations in order to recognize distinct signals. In contrast, the quality of fluorescence imaging data typically improves as the number of fluorescent molecules increases. Thus, appropriate conditions for both FCS and high quality fluorescence imaging are not wholly compatible with each other. Performing FCS in live cells presents additional hurdles because adjusting the density of fluorescent probes typically necessitates the regulation of protein expression.

In initial experiments performing FCS in live cells for measuring nuclear material properties, we expressed NLS-GFP-NLS in *S. pombe* cells (46). Although the expression level was appropriate for live cell imaging, the concentration of GFP in the nucleus in these experiments was too high to detect intensity fluctuations. This resulted in an autocorrelation curve characterized by low signal-to-noise ratio, which precluded accurate fitting for the determination of diffusion coefficients (Figure 1A). We addressed the overexpression by photobleaching the NLS-GFP-NLS to effectively reduce the concentration of fluorescent molecules. While this approach was somewhat effective (46), we found that it was technically difficult to reproducibly use photobleaching to tune fluorophore concentration, often resulting in poor fits (Figure 1B). Pre-bleaching also required us to use a higher laser power (relative to Figure 1A) for the single point FCS measurement, resulting in further bleaching during the FCS measurement. Additionally, deliberate photobleaching of cells prevents continued imaging following the FCS measurement, and could lead to off-target damage due to the creation of free radical oxygen species (48).

To address this challenge, we sought an alternative approach to effectively control the concentration of fluorescent molecules. The photoswitchable fluorophore mEos3.2 is ideal for this purpose because it is a fluorescent protein that photoswitches between two fluorescent states: one excited at 485 nm and one at 531 nm, with switching triggered by 405 nm illumination (Figure 1C, Supplementary Figure S1) (41). As a proof of principle, we activated a subset of molecules and thereby achieved sufficiently low concentrations of fluorescent molecules to effectively perform FCS and obtain autocorrelation curves that could be fit to extract diffusion parameters (Figure 1C).

Beyond activating only a subset of molecules, the photoswitching properties of mEos3.2 allow us to build an ideal workflow for both live cell imaging and FCS (Figure 2). First, we image in one channel (Channel 2, green) to monitor cellular morphology prior to activating a subset of fluorophores (Figure 2A). This ensures that the second channel (Channel 1, red) maintains a fluorophore concentration optimized for FCS measurements. Since photoswitching (Figure 2B) does not destroy the non-activated molecules, we can image in the appropriate channel before and after switching (Figure 2C). After photoswitching fluorophores, a single point measurement is collected by parking the confocal beam in the nucleus and measuring the intensity signal of diffusing fluorophores (Figure 2D, E). An intensity trace is produced from binning photon arrival macrotimes relative to the start of the experiment (Figure 2F) and the signal is autocorrelated using Eq. 1. After the autocorrelation curve is generated, the data is fit to a confocal diffusion model with an effective 3D observation volume using Eq. 2 (Figure 2G), which is consistent with a small molecule undergoing Brownian motion. From the fit, we extract physical parameters such as the average number of molecules in the confocal volume (*N*), and translational diffusion time (τ_D_), which we use as a metric for material properties of the nucleoplasm (Figure 2H). The diffusion coefficient (*D_1_*) is calculated from τ_D_ using Eq. 3.

**Figure 2:**
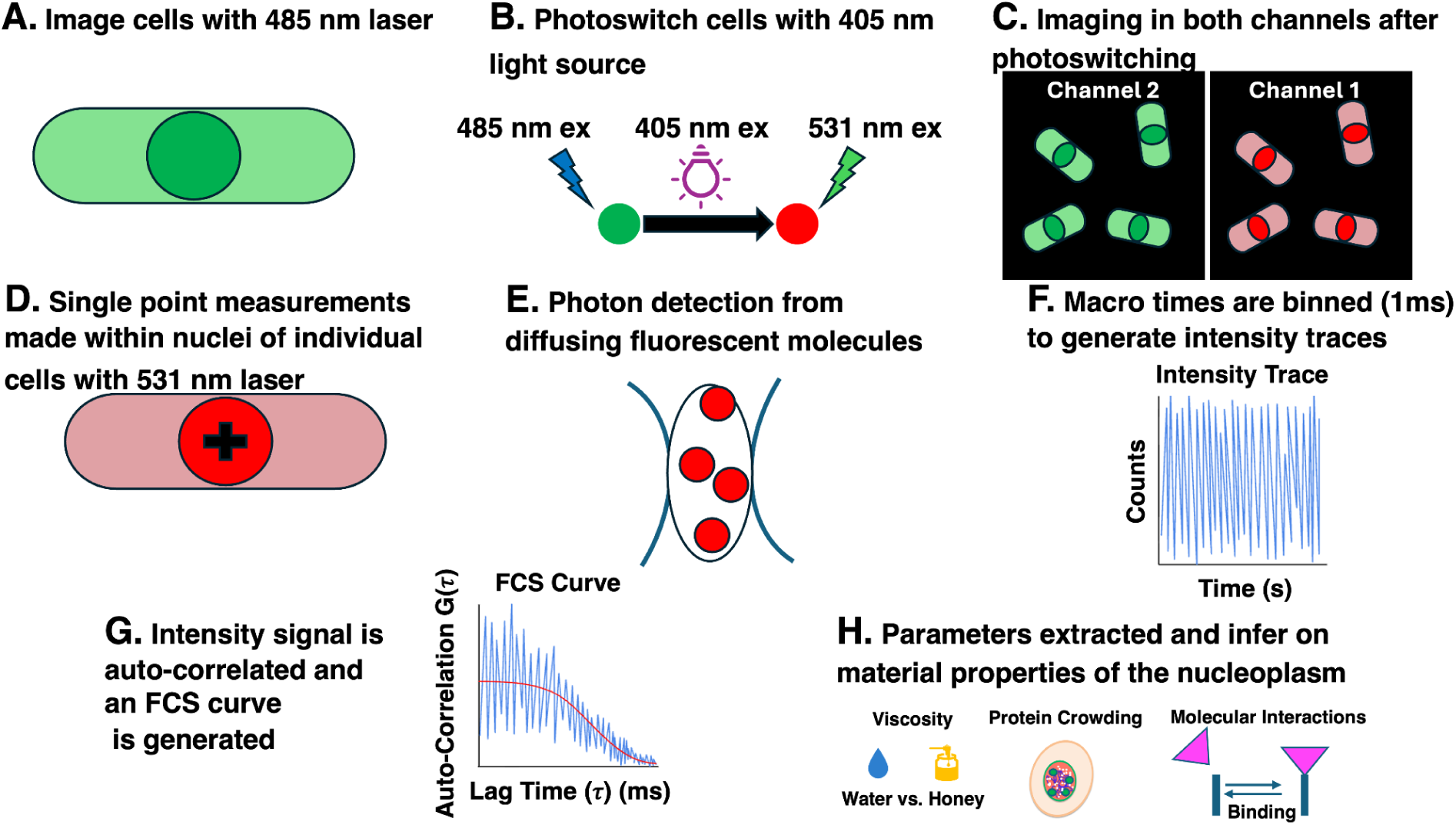
Workflow of FCS experiment in *S. pombe* cells expressing mEos3.2-NLS. (**A**) Cells are imaged with a 485 nm laser. (**B**) mEos3.2 fluorophores are excited with a 405 nm LED to photoswitch and then excited with a 531 nm laser to visualize fluorophores that switched fluorescent states in the cells. (**C**) Cells are visible in both spectrally separated channels before and after photoswitching. (**D)** Single point FCS measurements collected within nuclei of individual cells with a 531 nm laser. (**E**) Photons are detected as fluorophores diffuse through the confocal volume. (**F**) Photon macrotimes are binned (1 ms) to generate intensity traces. (**G**) Intensity signal is autocorrelated and an FCS Curve is generated. (**H**) Parameters and useful information such as diffusion coefficients and sample concentrations are extracted to gain insight into nucleoplasm material properties.

To precisely control the concentration of fluorescing molecules in the confocal volume, we next tuned the power or duration of the exposure to 405 nm LED activation light (Figure 3A, B). We confirmed that this approach allowed us to control the intensity of mEos3.2 fluorescence by quantitatively measuring photon counts in each channel as the power or duration of 405 nm exposure increased (Figure 3C). These experiments demonstrated that we could achieve appropriate intensity for FCS by controlling the 405 nm LED light exposure and/or duration, allowing us to estimate diffusion parameters from the fit (Individual intensity traces, FCS curves, and fit parameters are found in Supplementary Table S2 and Supplementary Figures S2, S3).

### Fluorescence Lifetime Imaging Microscopy (FLIM) verifies detection of mEos3.2

Autofluorescence from cellular materials often accompanies protein fluorescence. While standard live-cell imaging typically employs high expression levels to distinguish tagged molecules from this background signal, our approach deliberately uses low-intensity conditions. Consequently, verifying that intensity fluctuations originate from mEos3.2 rather than competing fluorescent species is essential. We address this by measuring the fluorescence lifetime—the average time a fluorophore remains excited before returning to the ground state—which serves as a molecular signature. With this information, we can create a FLIM image where the color of each pixel corresponds to a specific fluorescence lifetime (49) by creating a spatial map of average photon arrival times. In Figure 4A, we display a FLIM image of *S. pombe* MWE144 strain expressing the photoswitched state of mEos3.2-NLS. This is possible due to the Time-Correlated Single Photon Counting (TCSPC) (50), which simultaneously captures nanosecond-scale lifetimes and the millisecond-scale fluctuations required for FCS and FLIM measurements (Supplementary Figure S4(A)). By recording the delay between excitation and emission over multiple events, we construct a decay histogram from binning the nanotimes, and from binning the macrotimes we build an intensity trace for FCS (Supplementary Figure S4(B, C)). Fitting the excited state decay to a single exponential function yielded a lifetime of approximately 3.9 ns (Figure 4B), confirming the detection of mEos3.2. This lifetime is similar to 3.6 ns, which was previously measured for mEos3.2 in the red emitting state (51).

**Figure 3:**
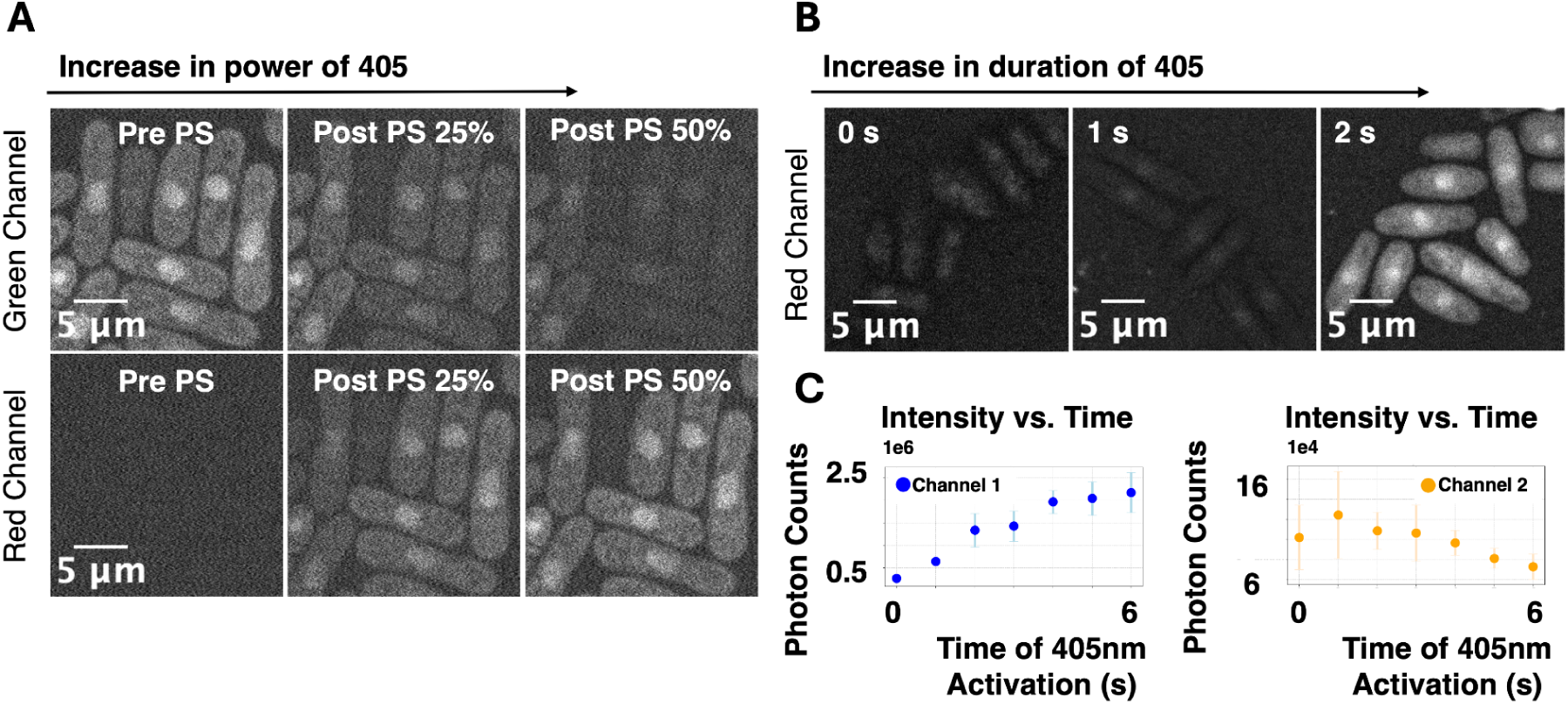
Tuning photoactivation time and power of 405 nm illumination. **(A)** *S. pombe* MWE144 strain expressing mEos3.2-NLS. Green channel shows cells before photoswitching (PS) with 405 nm LED light and then post-photoswitching at 25% and 50% power. Red channel shows cells after photoswitching with 405 nm LED light and then post-photoswitching at 25% and 50% power. Scale bar is 5 micrometers. **(B)** Representative images in the red channel (photoswitched state) show how the quality of images improve after prolonged 405 nm illumination. **(C)** Scatter plots showing increase in Channel 1 (red) photon counts and decrease in Channel 2 (green) photon counts as time of 405 illumination increases, indicating that fluorophores switch fluorescent states, corresponding to images in Figure 3B. Images in (A) obtained from Andor Dragonfly Spinning Disk Confocal and images in (B) obtained from Picoquant MicroTime200 Confocal Laser Scanning Microscope. Scale bar is 5 micrometers.

**Figure 4:**
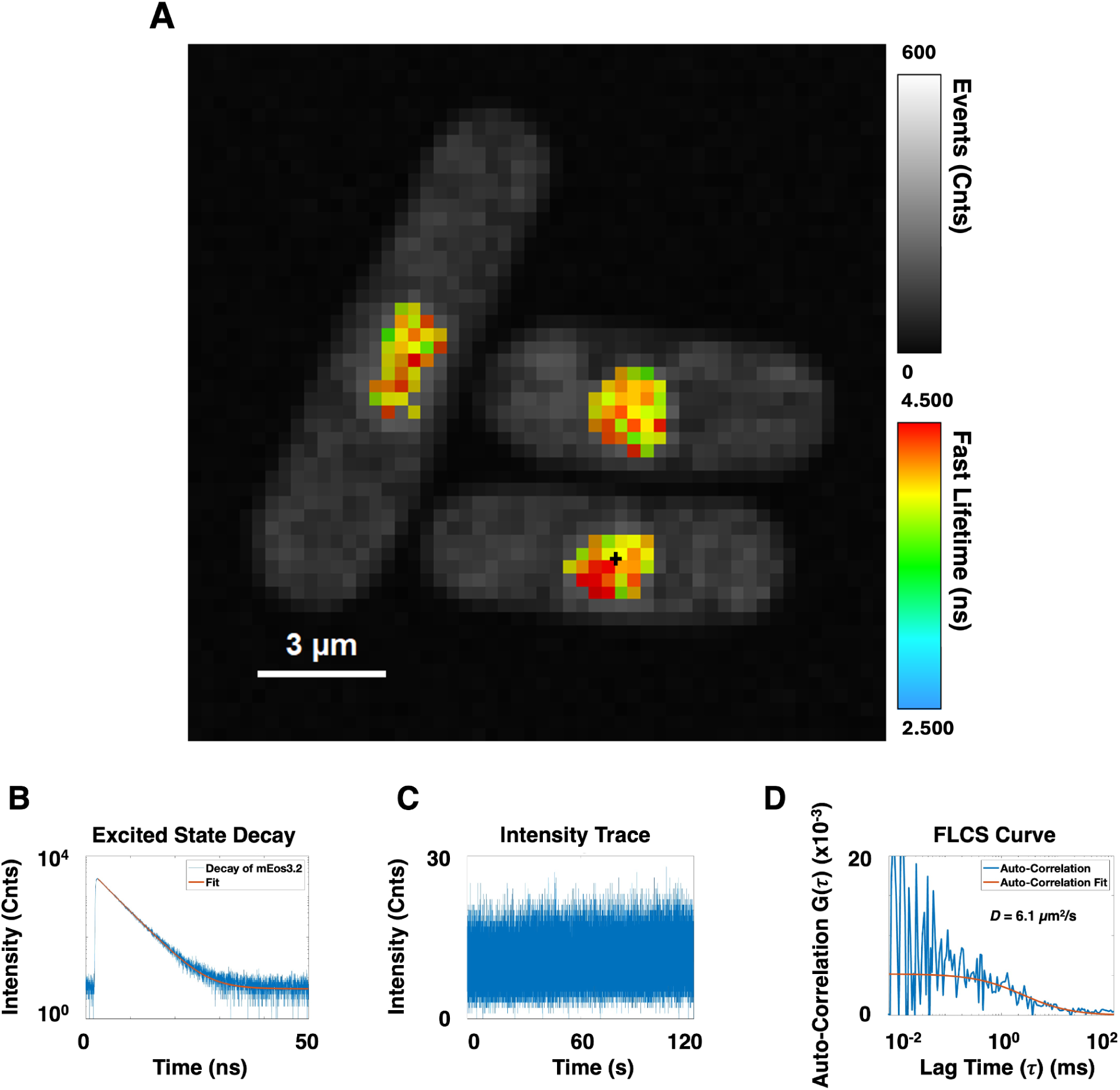
FLIM and FLCS of mEos3.2-NLS in *S. pombe* MWE144 strain **(A)** Fast Lifetime FLIM image of *S. pombe* MWE144 strain expressing mEos3.2-NLS, imaged in its photoswitched state, with colormap indicating fluorescence lifetime, calculated as described in Methods, with corresponding lifetime and intensity histogram shown in Supplementary Figure S5A and S5B. **(B)** Excited state decay of mEos3.2 showing the fluorescence lifetime extracted from fitting the photon data collected from FLCS analysis of photons collected at the location of the black cross in panel A (τ_1_ = 3.930 ns). **(C)** Intensity trace of single point FCS measurement collected in the cell pictured in panel A (black cross represents location of single point FCS measurement). **(D)** Autocorrelation and fit from FLCS analysis with a measured diffusion coefficient of 6.1 μm^2^/s.

Building upon our use of FLIM to confirm probe identity, we next considered how fluorescence lifetime information could be further integrated to improve our intensity based diffusion measurements. Fluorescence Lifetime Correlation Spectroscopy (FLCS) offers a refinement to traditional FCS by using the fluorescence lifetime of mEos3.2 as a filtering parameter prior to autocorrelation analysis. The filtering step is a mathematical function that weights the detected photon against its lifetime (Supplementary Figure S5(C)). A photon is weighted heavier if it matches the decay pattern of the fluorophore and less if not (52, 53). This approach allows us to selectively include photons originating specifically from mEos3.2 from the intensity trace (Figure 4C), thereby minimizing contributions from cellular autofluorescence or other background signals with distinct lifetime signatures when calculating the autocorrelation (Figure 4D). Because our setup employs TCSPC, it is already capable of capturing the lifetime information necessary for FLCS filtering. Incorporating FLCS improves the specificity and signal-to-noise ratio of FCS measurements, particularly in live-cell environments where background fluorescence or probe heterogeneity may otherwise confound interpretation. This methodological enhancement has the potential to yield more accurate diffusion coefficients and to better resolve subtle differences in nuclear material properties across cell cycle stages. Our measurements of diffusion coefficient by FCS and FLCS largely agree with each other (Figure 4D, Supplementary Figure S6(A-C)) indicating that even at the low fraction of photoswitched fluorophores we use, our measurements are still dominated by mEos3.2 molecules rather than autofluorescence.

### Single point Fluorescence Correlation Spectroscopy (FCS) and Engineered mEos3.2-NLS strain allow for visualization of the nucleus and detecting live cell dynamics

We can now use the combination of photoswitching and FCS to gain insight into nucleoplasm mechanics and its possible role in cell division. We hypothesized that differences in molecular diffusion may result from changes in physical organization of nuclear contents during cell division (54). To do so, we compare diffusion coefficients for interphase and mitotic cells (Figure 5).

**Figure 5:**
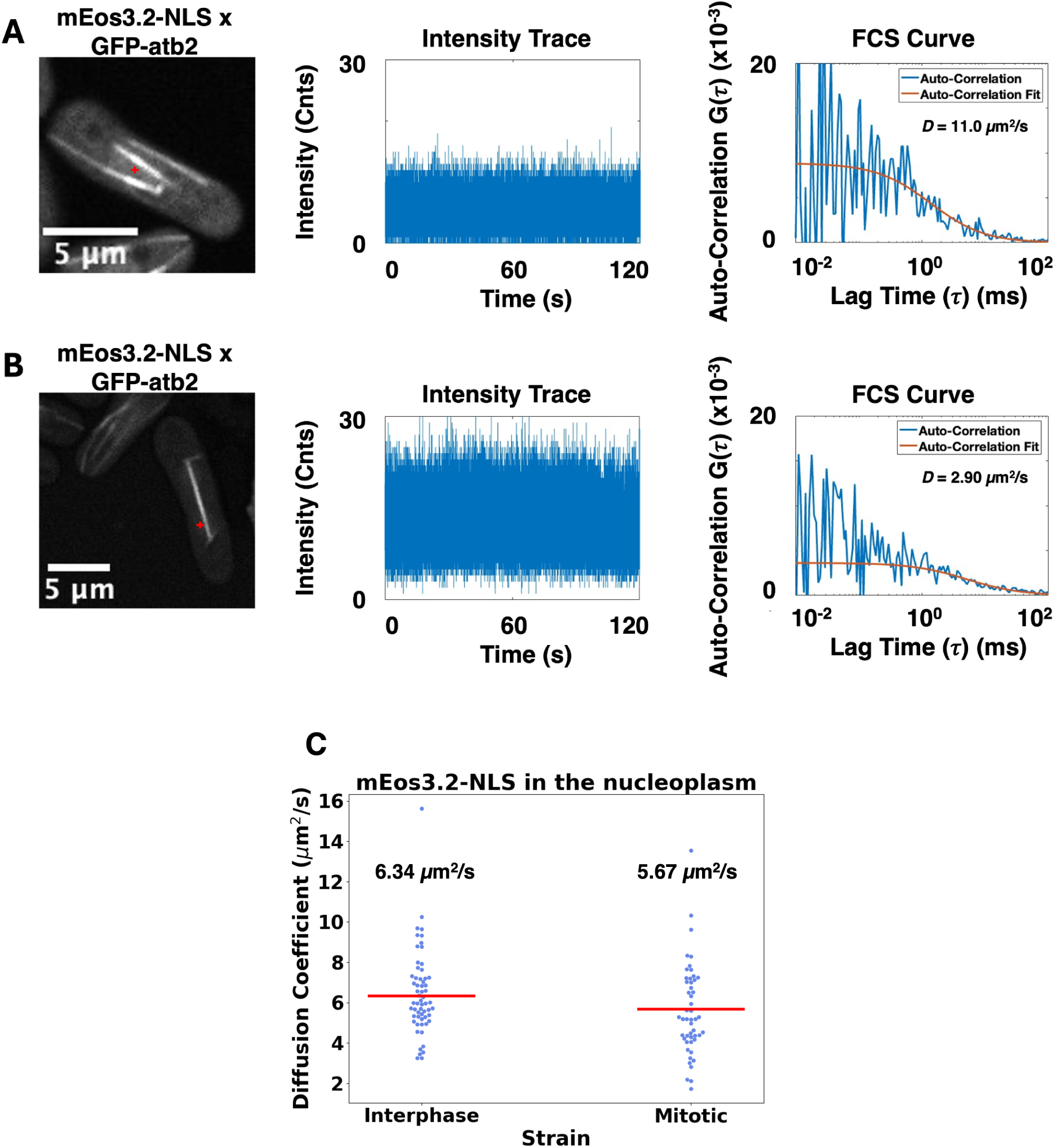
FCS data of *S. pombe* MWE145 strain expressing mEos3.2-NLS and GFP-Atb2. **(A)** Single point FCS measurement (red cross) within the nucleus of an interphase cell and its corresponding intensity trace, autocorrelation curve and fit with a measured diffusion coefficient of 11.0 μm^2^/s. Supplementary Figure S6(B) shows FCS vs. FLCS curves for interphase cell in Figure 5A. **(B)** Single point FCS measurement (red cross) within the nucleus of a mitotic cell and its corresponding intensity trace, autocorrelation curve and fit with a measured diffusion coefficient of 2.90 μm^2^/s. Supplementary Figure S6(C) shows FCS vs. FLCS curves for the mitotic cell in Figure 5B. Images are from green channel (native mEos3.2) and intensity traces and autocorrelations are in the red channel (photoswitched mEos3.2). **(C)** Comparing diffusion coefficients between interphase and mitotic cells. Blue dots represent individual cells and red bars represent average diffusion coefficients (N = 60 for interphase cells and N = 50 for mitotic cells). After measuring 60 interphase cells and 50 mitotic cells we measured an average diffusion coefficient of 6.34 μm^2^/s and 5.67 μm^2^/s, respectively. Welch’s t-test is performed for statistical analysis with a p-value of 0.08, indicating no statistically significant difference.

We performed single point FCS measurements in both non-dividing (interphase, Figure 5A) and dividing (mitotic, Figure 5B) cells and used them to calculate diffusion coefficients (Figure 5C). These data are consistent with globally similar diffusion rates between interphase and mitosis (Figure 5C), suggesting that, at the length and time scales probed by mEos3.2-NLS, nucleoplasm material properties do not undergo large changes across the cell cycle.

## Discussion

This study contributes to a growing body of work aimed at elucidating the material properties of intracellular compartments by leveraging diffusion as a functional readout (2, 20, 46). While previous research has demonstrated the utility of large fluorescent particles like GEMs in highlighting osmotic equilibrium and differences between nucleoplasmic and cytoplasmic environments (20), our approach takes it a step further by addressing some of the limitations inherent in using large molecular probes. In particular, GEMs—due to their size—may be excluded from certain subcellular regions, limiting the resolution and accuracy of diffusion-based measurements in structurally complex environments like the nucleus, which may exhibit poroelastic properties (54, 55).

To overcome these constraints, we introduced a small photoswitchable nuclear reporter, enabling diffusion measurements in regions that are difficult to access. In the future, this approach will allow for more nuanced insights into local viscosity, macromolecular crowding, and spatial heterogeneity within the nucleoplasm. Furthermore, our use of Fluorescence Correlation Spectroscopy (FCS), combined with fluorescence lifetime measurements (FLIM), offers a powerful means to simultaneously probe molecular mobility and environmental properties with high temporal resolution. While FCS alone is limited by concentration and spatial resolution constraints, we demonstrate that when paired with controlled photoactivation and time-resolved imaging, these limitations can be partially circumvented. This hybrid approach allows us to track the diffusion of small molecules in living cells with sufficient temporal and spatial resolution to extract meaningful material properties.

Diffusion influences key biological processes, such as nuclear transport, transcription regulation, and chromatin organization. Interestingly, our data indicate that diffusion rates for our nuclear reporter are globally similar between interphase and mitosis, despite vastly different nuclear organization. Furthermore, our measurements of nuclear diffusion using mEos3.2 are broadly consistent with the expected size dependence relative to prior diffusion measurements of 40 nm GEMs in S. pombe nuclei (20), suggesting that both probes report on a common underlying mechanical environment, despite differing by roughly an order of magnitude in size. These observations are consistent with the possibility that cells actively regulate their internal physical environment across the cell cycle. Together with our prior work (46) demonstrating strong mechanical coupling between the mitotic spindle and nuclear envelope during closed mitosis in *S. pombe*, our findings support the view that mitosis as a process is highly mechanically regulated.

Finally, the ability to simultaneously control probe concentration and measure diffusion through advanced fluorescence techniques opens up new avenues for studying intracellular dynamics. By fine-tuning molecular probe design and combining complementary imaging modalities, we can move toward a more comprehensive, spatially resolved map of intracellular material properties. This is relevant for understanding how perturbations in nuclear mechanics may contribute to pathological states or interfere with critical processes such as chromosome segregation. It is also important for clarifying how material properties vary across the cytoplasm, especially in light of recent evidence that processes such as biomolecular condensation can locally regulate diffusion even in the absence of membrane-enclosed compartments (55). Future studies should aim to further refine probe size and photophysical properties, improve multiplexing capabilities, and apply these techniques across different cell types and stress conditions. Integrating these data with quantitative measurements and complementary image correlation spectroscopy techniques such as RICS (56) will ultimately provide a more holistic understanding of cellular biophysics.

## Acknowledgements

We thank F. Chang and G. Dey for *S. pombe* strains and members of the Elting and LeBlanc labs for suggestions and advice. We thank the Wang Lab (NCSU) for sharing lab space and equipment. For microscopy support, we thank the Cellular and Molecular Imaging Facility (CMIF) at NCSU, which is supported by the State of North Carolina and the National Science Foundation. This work was supported by NIH 1R35GM138083 (MWE), NSF 2133276 (MWE), NIH/NCSU Molecular Biotechnology Training Program (MBTP) T32GM133366 (NZG), a Genetics and Genomics Scholarship (GGS) from NC State (NZG) and NC State Startup Funds and Chan Zuckerberg BioHub Science Diversity Leadership Award (SJL).

## Data Availability

Raw data is available upon request.

