## Supplementary Data for "Expression of photoactivatable molecules enables FCS in live cells by controlling fluorescence intensity"

### <u>Supplementary Figures and Tables</u>

| **Strain/protein** | **Genotype/sequence** |
|---|---|
| MWE2 | h+ GFP-atb2:kanMX ade6- leu1-32 ura4-D18<br>(Original source: Gift of Fred Chang, FC2861) |
| MWE38 | nup60-mCherry:Kan pBIP1-NLS-GFP-NLS:leu1+ ade- leu1-32 ura4D-18 h-<br>(Original source: Gift of Gautam Dey, GD250) |
| MWE41 | nem1::Hph nup60-mCherry:Kan<br>pBIP1-NLS-GFP-NLS:leu1+ ade- leu1-32 ura4D-18 h-<br>(Original source: Gift of Gautam Dey, GD258) |
| MWE53 | *S. pombe* h+ leu1-32 ura4 his7 lys1<br>(Original source: NBRP FY7455) |
| MWE133 | GFP-atb2:kanMX nem1::Hph<br>(Original source: this work produced by crossing MWE41 and MWE2 |
| MWE144 | his5::-mEos3.2-NLSSV40-natMX h+ leu1-32 ura4 his7 lys1<br>(Original source: this work, produced by transformation into MWE53) |
| MWE145 | his5:mEos3.2-NLSSV40-natMX,<br>GFP-atb2-kanMXleu1-32 ura4 his7 lys1<br>(Original source: this work, produced by crossing MWE144 and MWE133) |
| mEos-3.2 (codon optimized for *S. pombe*) | ATGTCAGCCATTAAGCCTGATATGAAGATCAAGTTAAG<br>AATGGAAGGTAATGTTAACGGACATCATTTCGTTATTG<br>ATGGAGATGGTACTGGTAAACCATTTGAAGGTAAACAA<br>TCAATGGATTTGGAAGTAAAGGAAGGTGGACCTTTACC<br>TTTCGCATTTGATATTCTTACTACTGCATTTCATTATG<br>GAAATAGAGTATTTGCAAAGTATCCAGACAATATCCAG<br>GATTACTTCAAGCAAAGTTTCCCTAAAGGTTACAGTTG<br>GGAACGATCTTTAACCTTTGAAGATGGAGGAATTTGCA<br>ATGCTAGAAATGACATTACCATGGAAGGTGATACCTTC<br>TATAACAAGGTTCGTTTCTATGGAACAAACTTTCCTGC<br>TAACGGACCTGTAATGCAGAAGAAGACTTTGAAATGGG<br>AACCTTCTACTGAGAAGATGTATGTTCGTGATGGAGTA |

| | |
|---|---|
| | TTGACAGGCGATATTGAAATGGCCTTATTGCTTGAGGG<br>AAACGCTCATTATCGTTGCGATTTCCGTACAACATACA<br>AAGCTAAAGAGAAAGGCGTAAAGTTGCCTGGTGCTCAC<br>TTTGTTGACCATTGCATAGAGATCCTTAGTCACGATAA<br>AGACTATAACAAGGTAAAGTTGTACGAACATGCTGTAG<br>CTCATTCAGGCTTACCCGATAATGCTCGACGA |

**Supplementary Table S1:** *S. pombe* strains in this work and their genotypes along with protein of interest, mEos3.2, and its codon optimized DNA sequence.

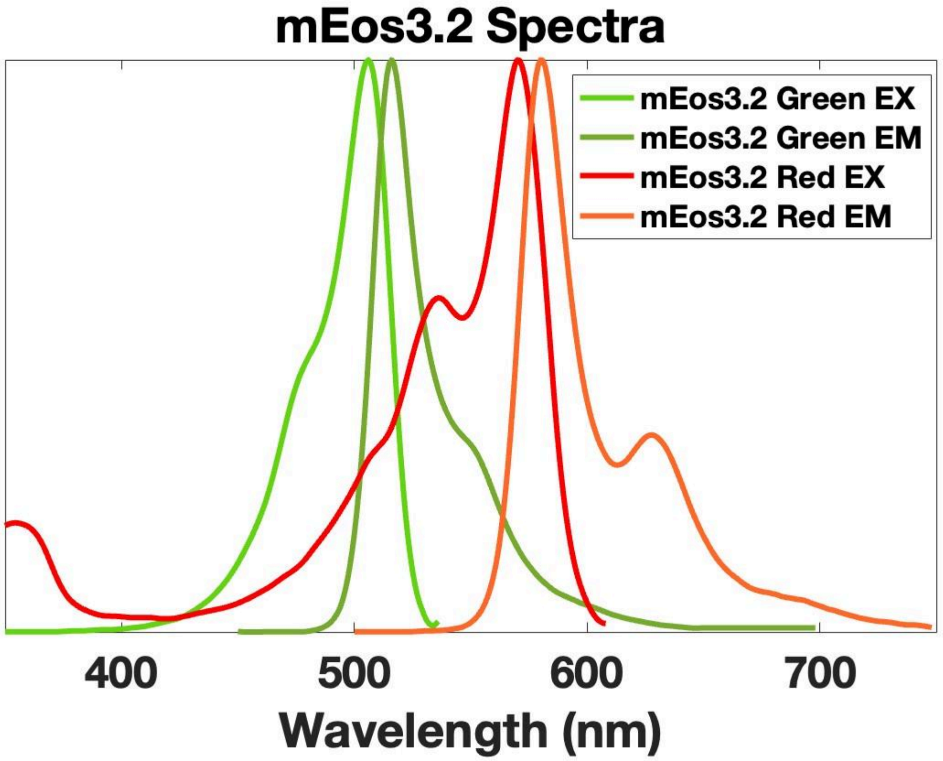


**Supplementary Figure S1:** Spectrum of mEos3.2 in both fluorescent states. Switching from green (first spectra) to red (second spectra) is induced from 405 nm illumination.

| Time of 405nm Illumination (s) | $\tau_{Diff}$ (ms) | N | $D_1$ ($\mu m^2/s$) | $\chi^2$ | $V_{eff}$ | $\kappa$ |
|---|---|---|---|---|---|---|
| 0 | 7 | 334 | 4 | 5.444 | 1.54 | 7.1 |
| 1 | 4.6 | 111 | 6.3 | 1.757 | 1.54 | 7.1 |
| 2 | 4.1 | 229 | 7 | 2.303 | 1.54 | 7.1 |
| 3 | 3 | 209 | 8.7 | 0.958 | 1.54 | 7.1 |
| 4 | 3.9 | 155 | 7.3 | 2.248 | 1.54 | 7.1 |
| 5 | 4.1 | 169 | 7 | 2.6 | 1.54 | 7.1 |

| | | | | | | |
|---|---|---|---|---|---|---|
| 6 | 4.2 | 255 | 6.8 | 1.63 | 1.54 | 7.1 |

**Supplementary Table S2:** FCS parameters showing time of 405nm illumination, $\tau_{Diff}$ (average time molecules spend in confocal volume), $N$ (Number of molecules in confocal volume), $D_1$ (Diffusion coefficient), $\chi^2$ (chi-square, how well data fits the model), $V_{eff}$ (Effective volume), and κ (ratio of width and length of confocal volume). When photoswitching is not conducted (405 nm illumination time = 0), there is insufficient signal, leading to a poor fit (see also Supplementary Figure S3, and correspondingly poor $\chi^2$). After photoswitching for at least 1 s, sufficient molecules are present to perform an effective FCS measurement.

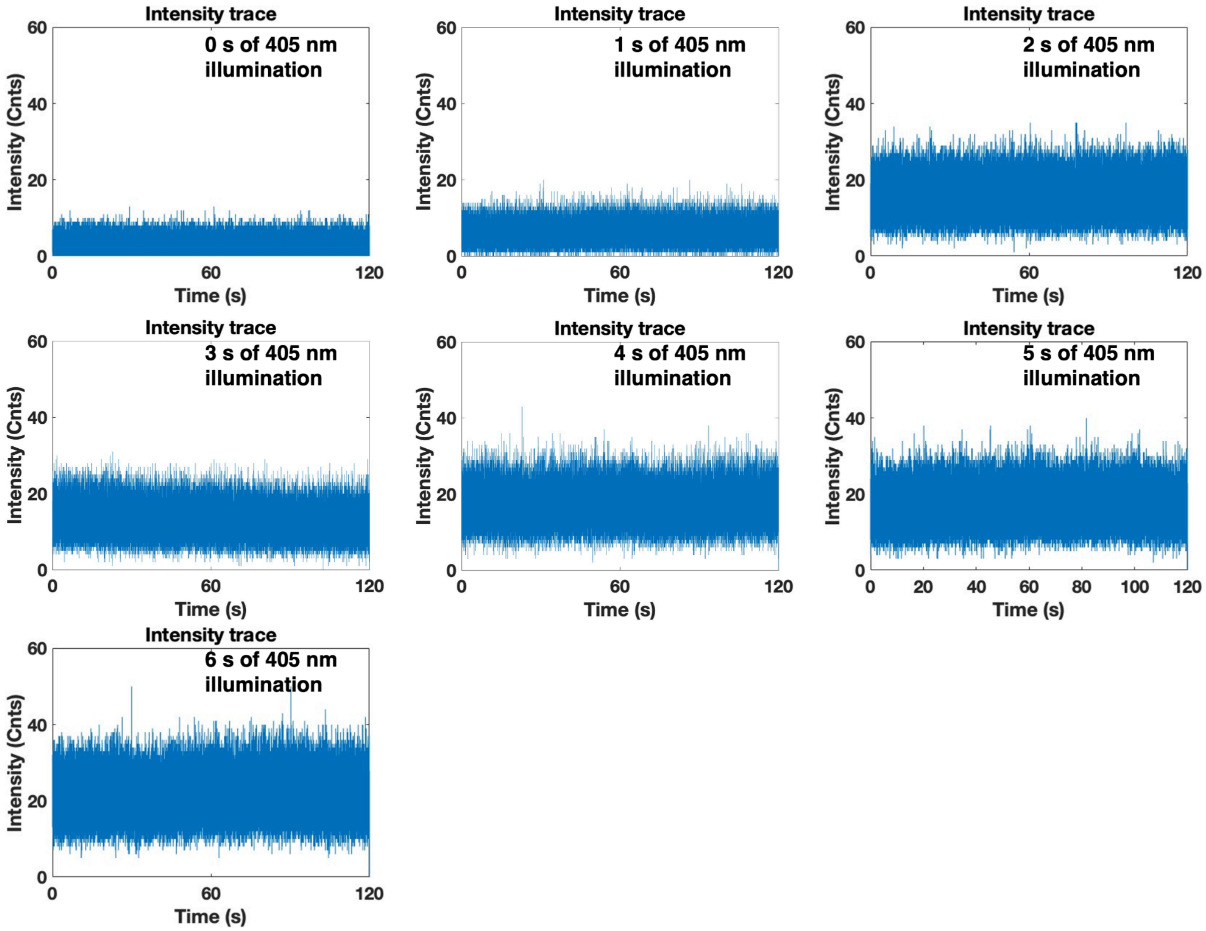


**Supplementary Figure S2:** Intensity traces from the red channel (photoswitched state) for a range of times from 0 to 6 seconds of 405nm illumination.

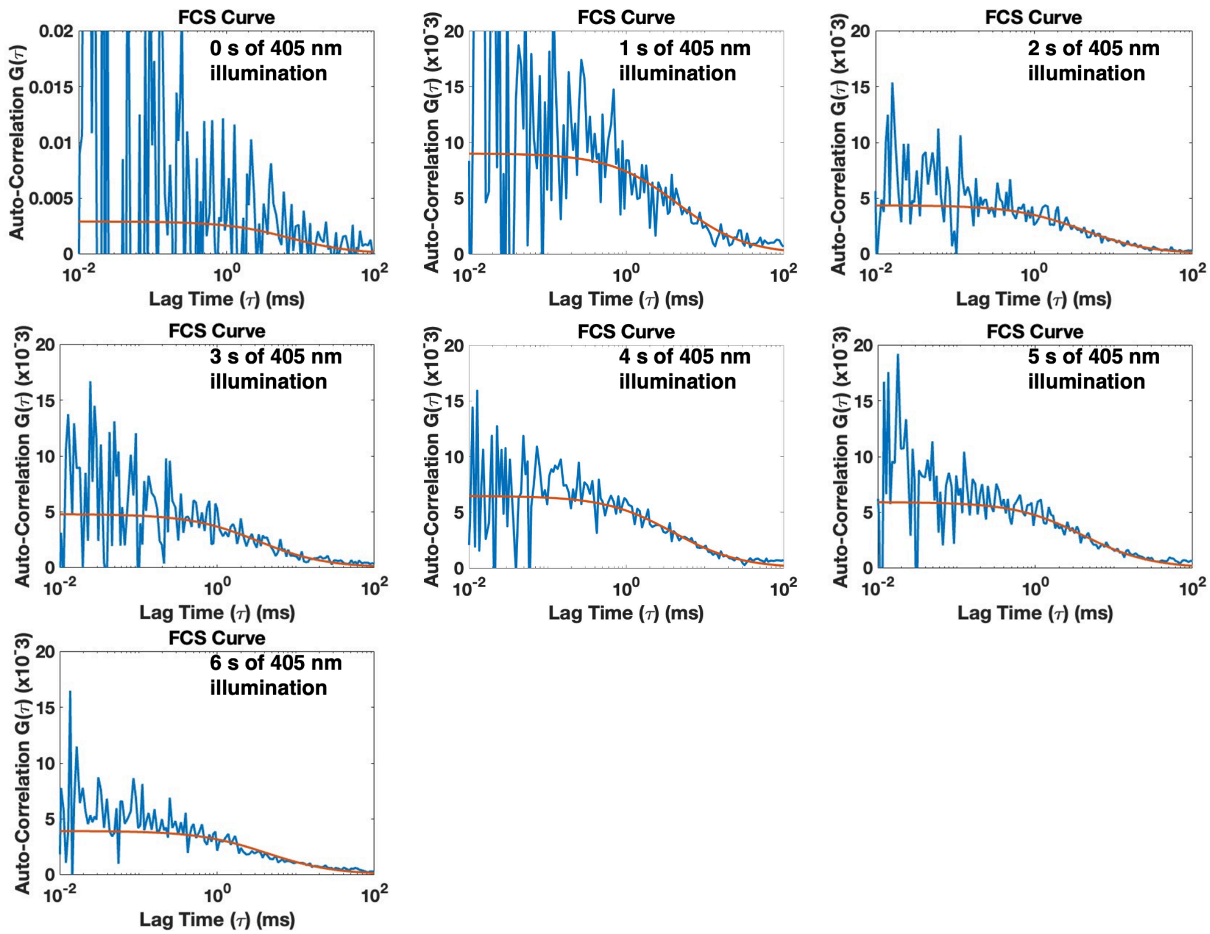


**Supplementary Figure S3:** Autocorrelation traces and fits from the red channel for a range of times from 0 to 6 secs of 405 nm illumination. The blue line is the autocorrelation curve and the red line is the autocorrelation fit.

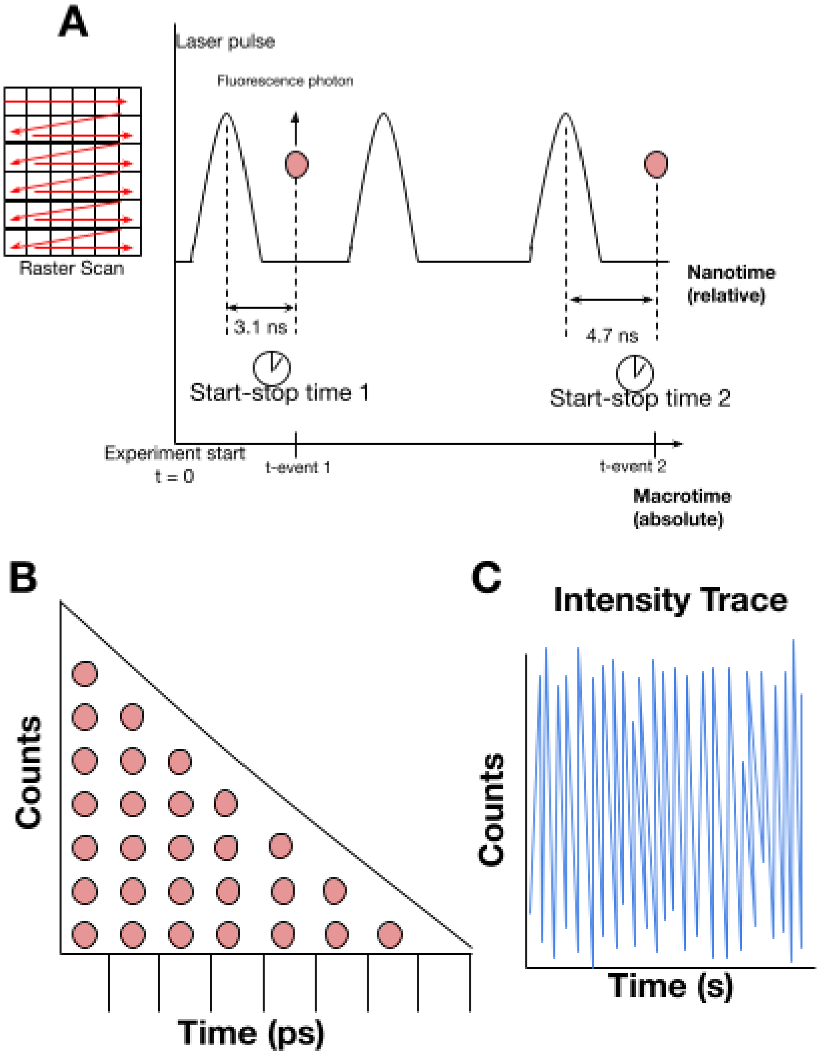


**Supplementary Figure S4: (A)** Principle of Time Correlated Single Photon Counting (TCSPC) with scanner and event timer (stop-watch). **(B)** Cartoon of binned photon nanotimes to create an excited state decay histogram. **(C)** Cartoon of intensity trace from binning of photon macrotimes.

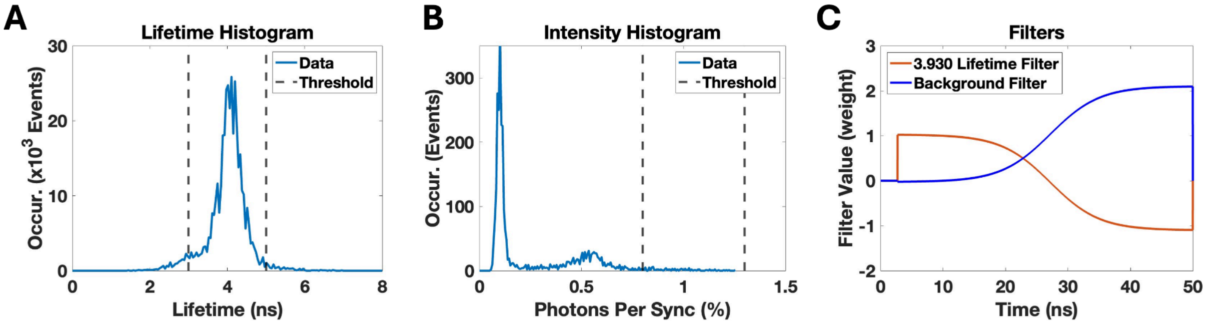


**Supplementary Figure S5**: Creation of FLIM image from Figure 4A **(A)** Lifetime histogram of fluorescent molecules in *S. pombe* MWE144 strain. **(B)** Intensity histogram showing the amount of photons detected per laser pulse and how many times that occurs. Black dashed lines on (A and B) represent thresholds for creating FLIM image in Figure 4A, which is discussed in the Methods section (main text). **(C)** FLCS Filters used for the autocorrelation of the signal from mEos3.2 based on its lifetime for Figure 4D.

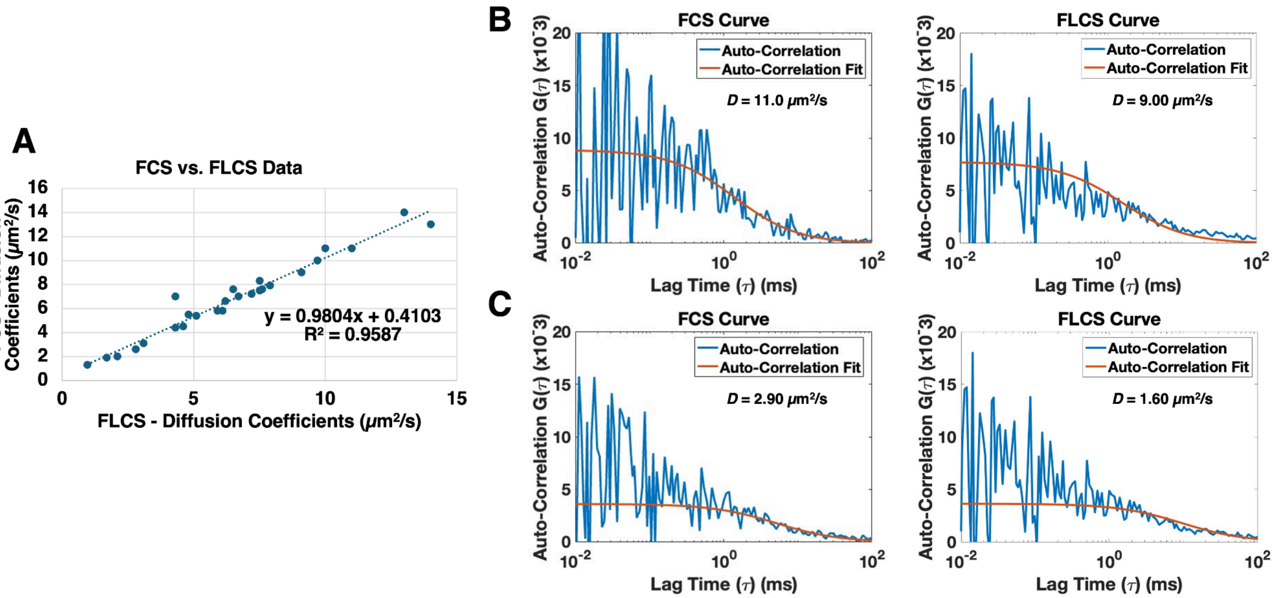


**Supplementary Figure S6**: (**A)** Comparing FCS and FLCS analysis of the same autocorrelation curves. Measurements were taken in *S. pombe* MWE144 strain expressing mEos3.2-NLS. (**B**) Comparing FCS vs. FLCS Curves from Figure 5A with a measured diffusion coefficient of 11.0 μm$^2$/s and 9.00 μm$^2$/s, respectively. (**C**) Comparing FCS vs. FLCS Curves from Figure 5B with a measured diffusion coefficient of 2.90 μm$^2$/s and 1.6 μm$^2$/s, respectively.
